# Evaluation of graphene oxide-mediated NET formation using HL-60-derived neutrophil-like cells

**DOI:** 10.1101/2025.06.06.658248

**Authors:** Kiyoshi Fukuhara, Takashi Obama, Hitomi Ohinata, Takashi Takaki, Masaru Kato, Rikako Ishigamori, Yukari Totsuka, Hiroyuki Itabe, Akiko Ohno

## Abstract

Neutrophil extracellular traps (NETs) are chromatin-based structures released by activated neutrophils in response to pathogens or chemical stimuli, contributing to host defense but also implicated in autoimmune disease and inflammation. As NET formation gains attention as an endpoint in in vitro immunotoxicity screening, the lack of reproducible and scalable systems hampers its broader application. Here, we developed an in vitro assay using HL-60-derived neutrophil-like cells (dHL-60) differentiated with all-trans retinoic acid to evaluate NET-inducing activity in a standardized, non-animal model. Graphene oxide (GO), a model nanomaterial known to trigger NETs in primary neutrophils, induced concentration-dependent NET formation in dHL-60 cells, with maximal induction at intermediate doses and attenuation at higher concentrations, likely due to particle aggregation. NET formation was validated by extracellular DNA staining and scanning electron microscopy. GO also induced superoxide-mediated ROS production, as confirmed by electron spin resonance, consistent with canonical NETosis pathways. Furthermore, GO suppressed PMA-induced NET formation, suggesting a dose- and context-dependent immunomodulatory effect. Collectively, our results demonstrate that dHL-60 cells recapitulate key features of NETosis observed in primary neutrophils and provide a practical, reproducible model for investigating immune responses to nanomaterials. This system supports the development of non-animal approaches for assessing immunological effects of chemical substances under controlled in vitro conditions.

## Introduction

The evaluation of chemical and pharmaceutical safety is a critical step in product development, traditionally relying on animal models to assess toxicity before clinical or commercial application. However, ethical concerns over animal welfare, coupled with increasing regulatory and societal pressure, have accelerated the pursuit of alternative testing strategies. In response, in vitro toxicity testing platforms are being actively developed, with a particular emphasis on generating systems that are mechanistically informative, reproducible, and biologically relevant. Among the major challenges in this field is the establishment of robust assays capable of detecting immune-related toxicities, especially those associated with neutrophil activation.

A promising endpoint in this context is the formation of neutrophil extracellular traps (NETs), web-like structures composed of decondensed chromatin and neutrophil-derived proteins that are released in response to microbial pathogens or sterile stimuli. [1] [2] [3] As a defense mechanism of the innate immune system, NETs contribute to microbial clearance and modulation of inflammation. However, aberrant or excessive NET formation has been increasingly associated with adverse health outcomes, including autoimmune diseases, thrombosis, and chronic inflammatory disorders. [4] [5] [6] Various exogenous agents, ranging from environmental pollutants [7] and pharmaceuticals [8] [9] to engineered nanomaterials, [10] [11] have been shown to induce NETosis, implicating it as a key event in the manifestation of immunotoxic effects.

The molecular process underlying NET formation involves a cascade of intracellular signaling events. Upon activation by pathogens or chemical stimuli, neutrophils generate reactive oxygen species (ROS) via NADPH oxidase and initiate chromatin decondensation through peptidylarginine deiminase 4 (PAD4)-mediated histone citrullination. [12] This leads to nuclear membrane disintegration, plasma membrane rupture, and the eventual extrusion of chromatin into the extracellular space. [13] While NETs serve a protective function, they also reflect an early immune activation state, rendering NET formation a potentially sensitive and mechanistically informative endpoint for in vitro toxicity testing, particularly in the context of immunotoxicity. [3] Despite this promise, conventional in vitro assays have largely focused on endpoints such as genotoxicity or cytotoxicity, which capture later-stage or downstream effects of toxicants. These methods often overlook early immune interactions, even though immune cells such as neutrophils are among the first to encounter xenobiotics after systemic exposure. Thus, assessing how neutrophils respond, especially through NET formation, provides a more physiologically relevant perspective on chemical bioactivity. However, primary human neutrophils are limited in vitro by their short lifespan (6–8 hours), donor-to-donor variability, and restricted scalability, posing significant barriers to assay standardization.

To address these limitations, HL-60 cells, a human promyelocytic leukemia cell line, can be differentiated into neutrophil-like cells (dHL-60) using all-trans retinoic acid (AtRA). [14] These cells exhibit essential neutrophil functions, including ROS generation and NET formation, and offer several practical advantages, such as consistent availability, reproducibility, and suitability for high-throughput applications. dHL-60 cells therefore represent a viable surrogate for primary neutrophils in NET-based toxicological assessments.

Nanomaterials such as graphene oxide (GO) have attracted considerable attention in biomedical and industrial contexts owing to their unique physicochemical properties. However, these same properties also complicate the prediction of biological outcomes. GO has been reported to induce NET formation in primary neutrophils, [15] [9] raising concerns regarding its potential immunomodulatory activity. In this study, we utilized GO as a model nanomaterial to validate an in vitro NET formation assay using dHL-60 cells. We demonstrate that GO elicits NET formation via canonical ROS-dependent pathways and modulates NET responses to known stimulants such as PMA.

By establishing that dHL-60 cells respond to GO with reproducible NET formation and capture complex modulatory effects, this study lays the foundation for a scalable, non-animal assay system targeting NETs as a mechanistic endpoint. Such systems may not only facilitate the mechanistic dissection of nanomaterial-immune interactions, but also support the development of predictive and ethically responsible platforms for immunotoxicity screening and chemical safety evaluation.

## Materials and methods

### Chemicals

AtRA and phorbol 12-myristate 13-acetate (PMA) were purchased from Fujifilm Wako Pure Chemical Co. (Osaka, Japan). SYTOX Green was purchased from Thermo Fisher Scientific (Rockford, IL, USA). Furthermore, 5,5-dimethyl-1-pyrroline N-oxide (DMPO) was purchased from Labotec (Tokyo, Japan). Graphene oxide (777676) and poly-L-lysine solution (P4707) were purchased from Sigma–Aldrich (Saint Louis, MO, USA). Graphene oxide (GO) was suspended in a serum-free Roswell Park Memorial Institute (RPMI)-1640 (phenol red free) medium at the appropriate concentration and underwent sonication at room temperature for 1 min immediately before each experimental procedure to ensure proper dispersion.

### NET induction by HL-60 cell-derived neutrophils

HL-60 cells were treated with AtRA to induce their differentiation into neutrophil-like cells, as previously described [16]. Briefly, 2 μM AtRA were added to HL-60 cells (2.0 × 10^6^ cells/dish) cultured in an RPMI-1640 medium supplemented with 5% deactivated (for 30 min at 56 □) fetal bovine serum, 50 U/mL penicillin, and 50 μg/mL streptomycin in 10 cm dishes for 4 days. HL-60-derived neutrophil-like cells were collected and washed once with serum-free RPMI-1640 (phenol red free) medium, seeded in a 24-well poly-l-lysine-coated plate (5.0 × 10^5^ cells/well), and cultured for 30 min. The cells were treated with aliquots of PMA or GO and incubated at 37 □ under 5% CO_2_ for 2 h.

### Fluorometric quantitation of NET-DNA

The culture medium was treated with 1 U/mL MNase to degrade the DNA exposed extracellularly upon NET formation in the culture dishes. The culture medium from each dish was recovered and centrifuged at 1,800 × *g* for 10 min. After mixing the supernatant with 1 μM SYTOX Green, the fluorescence intensity was measured as previously described [16]. Fluorescence (ex: 485 nm, em: 525 nm) was measured using Varioskan Flash (Thermo Fisher Scientific).

### Dynamic light scattering (DLS) analysis

DLS measurements were conducted using a Nanotrac Wave II (Microtrac Inc.) at room temperature. Before measurement, the nanoparticle suspension was prepared at the appropriate concentration in an RPMI medium and underwent sonication for 1 min at room temperature. The suspension was allowed to stand for 10 min before DLS analysis. The particle size distribution was analyzed based on the intensity-weighted average diameter.

### Electron spin resonance (ESR)-spin trapping ROS determination

HL-60-derived neutrophil-like cells were seeded in a 24-well plate coated with poly-L-lysine (5.0 × 10 □ cells/well) and incubated for 30 min at 37 □under 5% CO□. The cells were treated with DMPO at a final concentration of 400 mM, and PMA or GO was added. After incubation for 10 or 20 min at 37 □ under 5% CO □, the culture medium was transferred to a flat quartz ESR cell and placed in a JEOL JES-3X ESR spectrometer. The ESR spectra were recorded with manganous oxide as an external standard at room temperature under the following conditions: microwave power, 10.0 mW; modulation frequency, 100 kHz; modulation amplitude, 0.63 G; scanning field, 3,360 ± 50 G; receiver gain, 6.3 × 100; response time, 0.1 s; sweep time, 2 min.

### Scanning electron microscopy (SEM) analysis

HL-60 cells were differentiated into neutrophil-like cells through treatment with AtRA. The differentiated cells were seeded onto chamber slides (Cat. No. 154526PK; Thermo Fisher) and incubated for 30 min at 37□in a 5% CO□ atmosphere to ensure adherence. Subsequently, the cells were stimulated with 50 nM PMA or 100 µg/mL GO and incubated for 2 h under the same conditions. Following stimulation, the cells were fixed with 4% paraformaldehyde in phosphate-buffered saline for 15 min at room temperature. After fixation, samples were processed for SEM imaging.

### Statistical analysis

Data were expressed as mean ± SD. The comparisons were executed through one-way analysis of variance (ANOVA) followed by Dunnett’s post hoc test. Statistically significant difference was evaluated at p < 0.05.

## Results

### GO induces NET formation in dHL-60 cells

Phorbol 12-myristate 13-acetate (PMA) is a well-established chemical stimulant known to activate protein kinase C (PKC), thereby triggering neutrophil extracellular trap (NET) formation in various neutrophil models, including HL-60 cells differentiated with AtRA. [17] To explore whether graphene oxide (GO), previously reported to induce NETs in primary neutrophils, exerts a similar effect on dHL-60, we evaluated NET formation in response to GO stimulation.

GO was ultrasonically dispersed in culture medium to achieve uniform particle distribution and incubated with dHL-60 cells for 2 hours. The extent of NET formation was quantitatively assessed by SYTOX Green fluorescence, which selectively stains extracellular DNA released during NETosis (Figure 1A). GO stimulation led to significant DNA release in dHL-60 cells, confirming NET formation. Notably, treatment with 25 µg/mL GO elicited a stronger NETotic response than 50 nM PMA, a potent positive control. GO-induced NET formation exhibited a clear concentration-dependent increase, peaking at 50 µg/mL. However, at 100 µg/mL, a paradoxical reduction in NET formation was observed.

**Fig 1.**
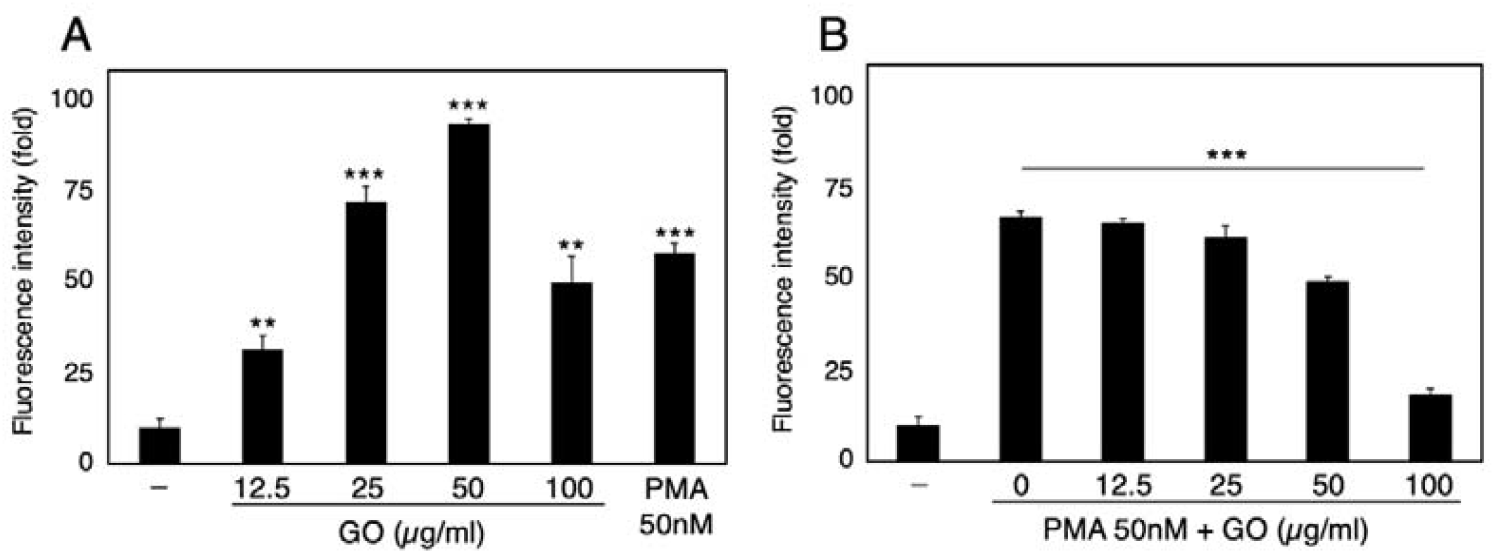
Fluorometric quantification of DNA release from HL-60-derived neutrophils **(A)**. Graphene oxide (GO; 12.5−100 µg/mL) and phorbol 12-myristate 13-acetate (PMA; 50 nM) were added to HL-60-derived neutrophils, and the mixture was incubated for 2 h. (B) HL-60-derived neutrophils were stimulated with 50 nM PMA for 10 min and treated with 0−100 µg/mL GO for an additional 2 h. The collected culture medium was treated with micrococcal nuclease and stained with 1 µM SYTOX Green, after which DNA released from the cells was quantified fluorometrically. Data are presented as the mean ± SD of four independent experiments. Asterisks indicate statistical significance. \*\**p* < 0.005, \*\*\**p* < 0.001.

This non-linear response suggests that higher GO concentrations may result in particle aggregation, thereby impairing effective interaction with the cell surface. Indeed, the bioactivity of nanomaterials is often dictated by their dispersion state, and loss of colloidal stability at high concentrations can reduce cellular uptake and membrane association. These findings underscore the importance of GO concentration and dispersion state in modulating NET-inducing capacity in vitro.

### GO modulates PMA-induced NET formation

PMA activates NETosis by engaging PKC at the plasma membrane, and its effect can be modulated by co-exposure to pro-oxidant or antioxidant compounds. Previous studies have shown that oxidized low-density lipoprotein (oxLDL) enhances, while resveratrol inhibits, PMA-induced NET formation. [16] [18] To determine whether GO influences NET formation indirectly through modulation of known stimuli, we examined the effect of co-treatment with GO and PMA in dHL-60 cells.

Stimulation with 50 nM PMA alone robustly induced NET formation. However, the addition of GO during PMA stimulation significantly suppressed NET formation in a concentration-dependent fashion (Figure 1B). At 100 µg/mL, GO exhibited the most pronounced inhibitory effect. These results suggest that GO may antagonize PKC-driven NETosis, potentially by disrupting redox-sensitive signaling pathways or membrane-associated processes. Such dual functionality, inducing NETs independently, while suppressing chemically triggered NETosis, highlights the complex, context-dependent bioactivity of GO.

### GO aggregation at high concentrations affects its biological activity

To further probe the concentration-dependent effects of GO on NET formation, we characterized its dispersion profile in cell culture medium RPMI via DLS (Figure 2). GO at 20 µg/mL exhibited a unimodal size distribution with a median hydrodynamic diameter (D50) of 838 nm, indicating well-dispersed nanosheets. As the concentration increased, a second peak appeared, reflecting the formation of larger particle aggregates. At 200 µg/mL, D50 increased markedly to 2518 nm, consistent with extensive aggregation.

**Fig 2.**
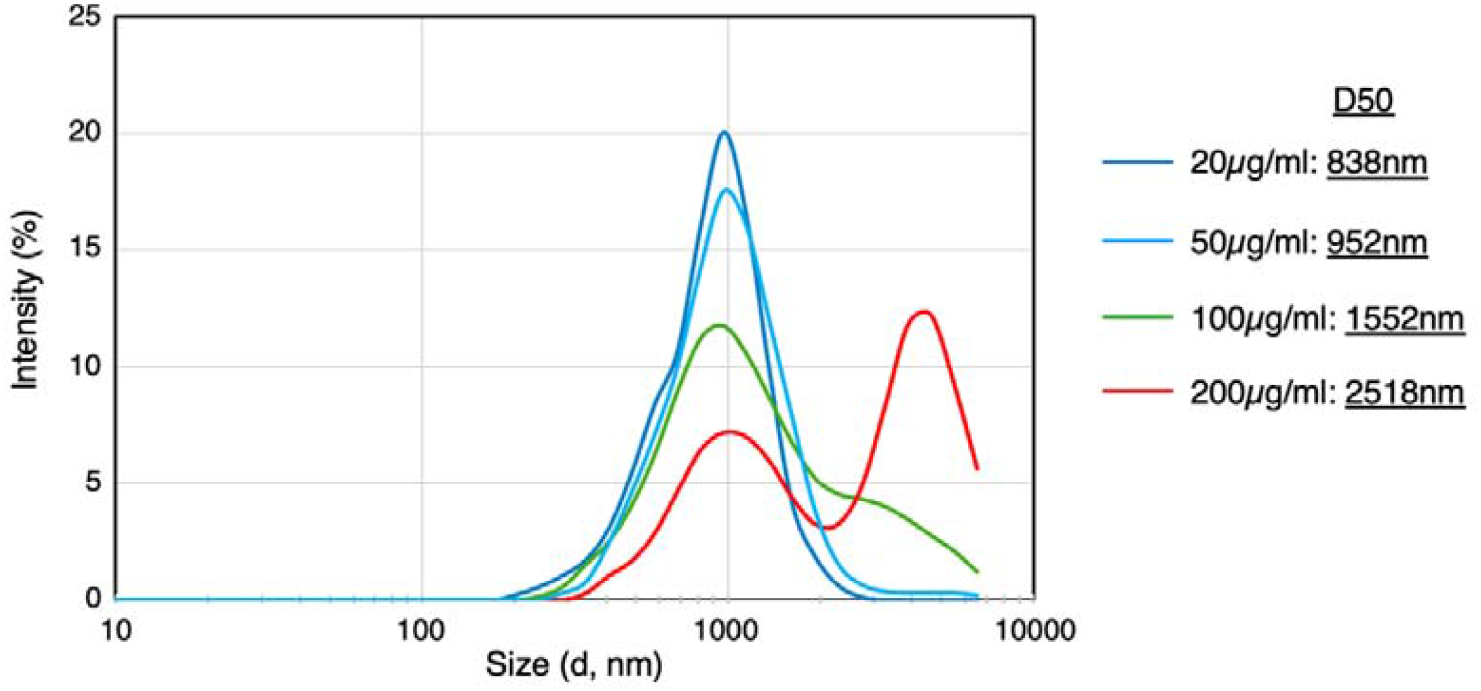
Dynamic light scattering (DLS) analysis of GO (20−200 µM) in RPMI.

This aggregation likely reduces GO’s surface area available for cellular interaction and may hinder its internalization or membrane binding. The loss of NET-inducing capacity at 100 µg/mL may therefore be attributed to this physical transformation. These results reinforce the concept that nanomaterial dispersion is a critical determinant of biological activity and must be carefully monitored in concentration-response evaluations.

### GO induces ROS production in dHL-60 cells

NET formation is tightly linked to intracellular ROS production, primarily driven by the NADPH oxidase complex during the respiratory burst. [19] To investigate whether GO induces ROS in dHL-60 cells, we performed ESR spectroscopy with the spin-trap 5,5-dimethyl-1-pyrroline N-oxide (DMPO) (Figure 3). dHL-60 cells stimulated with PMA displayed a characteristic ESR signal for the DMPO-OH adduct, a spin-trap product of hydroxyl radicals, with a 1:2:2:1 quartet hyperfine pattern (a_N = 1.49 mT, a_H = 1.49 mT). The signal was attenuated by superoxide dismutase (SOD), confirming that hydroxyl radicals originated via superoxide-dependent pathways.

**Fig 3.**
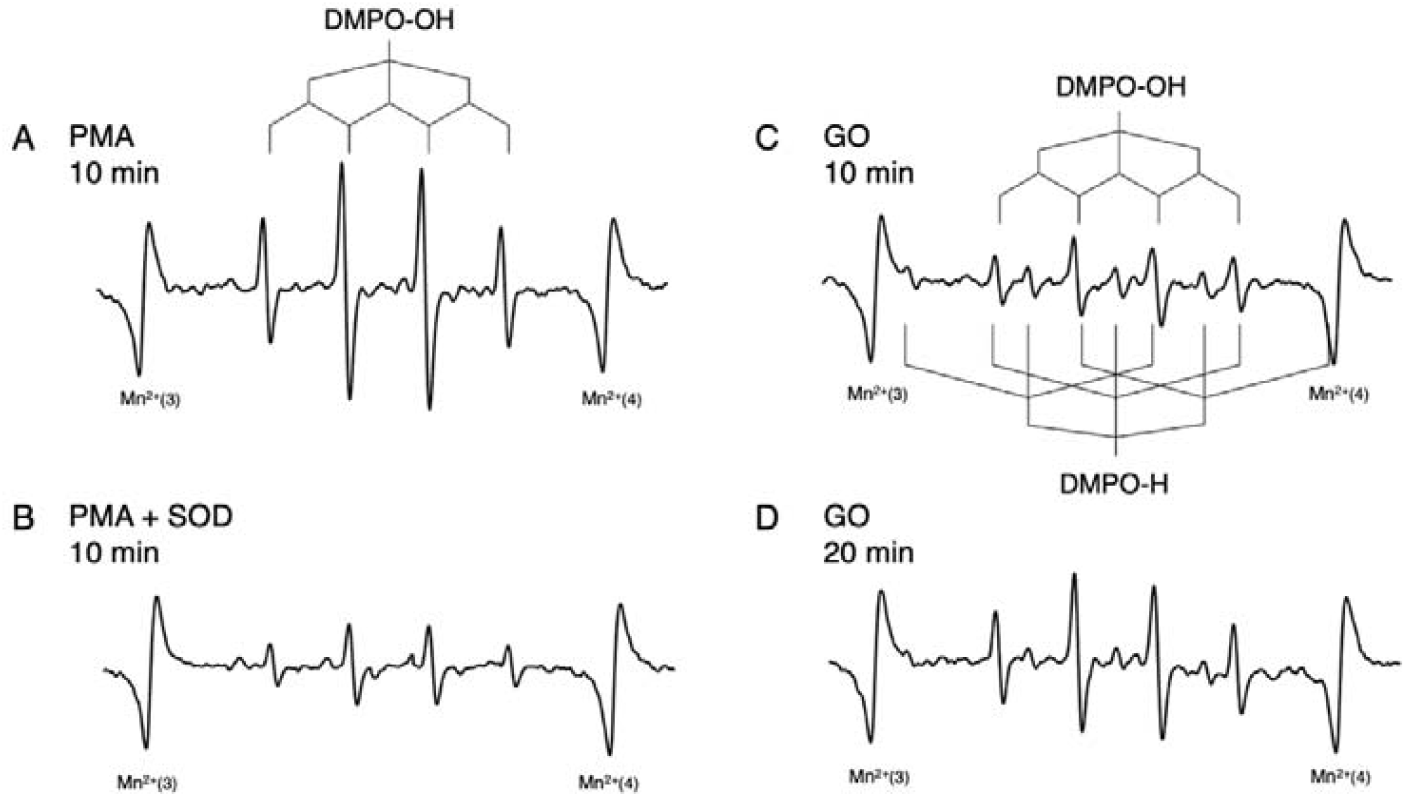
Electron spin resonance (ESR) spectra of 5,5-dimethyl-1-pyrroline N-oxide (DMPO)-free radical spin adducts obtained from solutions containing dHL-60 treated with PMA or GO. (A) 50 nM PMA + 613 mM DMPO, 10 min incubation. (B) Similar to that in (A), with the addition of 100 U/mL superoxide dismutase (SOD). (C) 100 µM GO + 613 mM DMPO, 10 min incubation. (D) Similar to that in (C); however, with 20 min incubation.

Similarly, GO stimulation yielded a strong DMPO-OH signal, supporting the conclusion that GO activates ROS production through a comparable mechanism. Additionally, a weaker triplet-of-triplets signal (1:1:2:1:2:1:2:1:1) characteristic of DMPO-H, the adduct of hydrogen radicals (aN = 1.63 mT, aH = 2.53 mT), was observed in GO-treated cells. While the biological implications of hydrogen radical formation remain unclear, this finding points to the distinct redox properties of GO and warrants further mechanistic investigation.

### Morphological validation of GO-induced NET structures

To confirm the morphological characteristics of GO-induced NET formation, we conducted SEM analysis of dHL-60 cells exposed to GO or PMA (Figure 4). In PMA-treated cells, extensive chromatin decondensation and extracellular web-like structures were observed, consistent with classical NETosis. GO-treated dHL-60 cells exhibited comparable phenotypes, including membrane rupture and the release of reticulated chromatin fibers.

**Fig 4.**
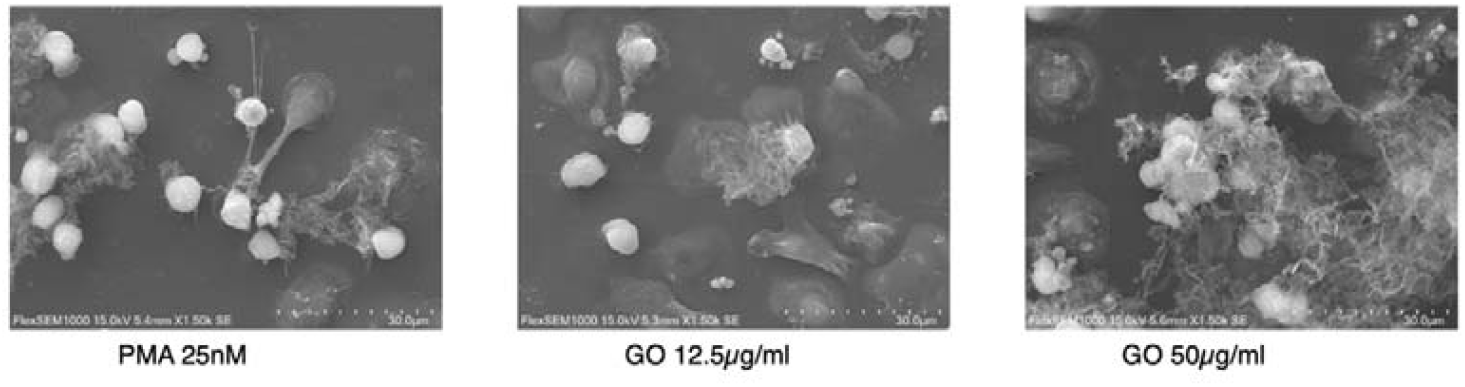
Scanning electron microscopy (SEM) images showing the characteristic appearance of NETs in dHL-60 cells treated with PMA or GO.

These ultrastructural findings provide visual evidence that GO induces NET formation via a process phenotypically indistinguishable from canonical NETosis. When considered alongside DNA quantification and ROS measurements, these results establish that GO elicits a bona fide NET response in dHL-60 cells, validating the use of this cell model for NET-oriented toxicity screening.

## Discussion

The biological impact of chemical substances and engineered nanomaterials extends beyond classical toxicological endpoints, increasingly encompassing immunological responses such as inflammation, immune suppression, and aberrant activation. Among these, the formation of NETs has emerged as a highly sensitive indicator of immune cell activation and dysregulation. Although originally identified as a host defense mechanism against microbial invasion, excessive or misregulated NET formation has been implicated in diverse pathologies, including autoimmune diseases, thrombosis, and tissue damage. [20] [6] These insights underscore the need to incorporate NET formation as a biologically relevant endpoint in next-generation in vitro toxicology platforms. However, the practical implementation of NET-based assays has been hampered by the technical limitations of primary neutrophils, such as their short lifespan, donor-to-donor variability, and restricted scalability. To circumvent these issues, we employed HL-60-derived neutrophil-like cells (dHL-60), which acquire key neutrophilic functions upon differentiation with AtRA. Our study demonstrates that dHL-60 cells robustly reproduce canonical features of NETosis, including extracellular DNA release, reactive oxygen species (ROS) generation, and characteristic chromatin morphology, in response to both phorbol ester stimulation and nanomaterial exposure.

GO was used as a model nanomaterial due to its reported ability to induce NET formation in primary neutrophils. Consistent with previous findings, GO elicited pronounced NET formation in dHL-60 cells in a concentration-dependent manner. The peak response observed at 50 µg/mL, followed by attenuation at higher concentrations, suggests that GO’s bioactivity is modulated by its dispersion state. DLS analysis revealed significant aggregation at elevated GO concentrations, which likely reduced its effective cellular interactions. This phenomenon highlights the critical role of nanomaterial physicochemical properties, particularly particle size and surface area, in determining biological activity. Similar behaviors have been reported for other nanomaterials, such as silica [21] and titanium dioxide, [22] indicating that this is a generalizable concern in nanotoxicology.

Importantly, our assay system also captured the modulatory effects of GO on chemically induced NETosis. Co-treatment with PMA and GO resulted in a significant, dose-dependent suppression of NET formation. This dual functionality, namely NET induction in naïve cells and inhibition of PMA-triggered NETosis, suggests that GO can interact with both basal and activated signaling pathways. While the underlying mechanisms remain to be fully elucidated, possible explanations include interference with PKC signaling, alteration of redox balance, or modulation of membrane-associated receptors. These results underscore the utility of the dHL-60 model not only for screening NET inducers, but also for identifying potential modulators or inhibitors of NETosis, which may have therapeutic relevance.

The involvement of ROS in NET formation is well established, with NADPH oxidase-mediated superoxide production serving as a critical upstream trigger. ESR spectroscopy revealed that GO induces hydroxyl radical formation via superoxide-dependent pathways, mirroring the ROS signature observed with PMA. Notably, GO also generated hydrogen radical signals, a phenomenon not typically reported in PMA-induced NETosis. Although the functional implications of hydrogen radical generation remain unclear, this observation suggests that GO may activate distinct redox chemistries, potentially contributing to its unique biological effects.

Morphological analyses via scanning electron microscopy provided further validation of NET-like structures induced by GO, confirming the rupture of nuclear and plasma membranes and the release of extracellular chromatin webs. Taken together with fluorescence-based DNA detection and ROS measurements, these findings establish that dHL-60 cells can serve as a robust and reproducible surrogate model for NET-based toxicity assessment.

Beyond the scope of GO, our previous unpublished data indicate that other particulate materials, such as silica nanoparticles, also induce NET formation in dHL-60 cells. These observations support the broader applicability of this model system across diverse compound classes. Given its reproducibility, scalability, and non-reliance on animal-derived primary cells, the dHL-60 NET assay is particularly suited for high-throughput screening and mechanistic investigations in immunotoxicology. Nevertheless, several limitations should be acknowledged. The mechanistic equivalence between NETosis in dHL-60 cells and primary neutrophils requires further validation, particularly with respect to downstream immunological consequences such as cytokine release, macrophage activation, or tissue remodeling. Additionally, while ESR spectroscopy offers valuable insights into ROS dynamics, complementary assays, including mitochondrial function, calcium flux, and transcriptional profiling, may enrich mechanistic interpretation. Finally, as nanomaterials display complex behaviors in biological fluids, standardization of dispersion protocols will be essential for assay reproducibility and inter-laboratory comparability. Collectively, these results support the use of HL-60-derived neutrophil-like cells as a robust and reproducible platform for dissecting NET-related cellular responses to chemical substances and nanomaterials. The ability of this system to detect both NET-inducing and NET-modulating activities, together with its compatibility with high-throughput formats, underscores its potential utility in mechanism-driven, non-animal immunotoxicity testing strategies.

## Conclusion

This study establishes a standardized and mechanistically informative in vitro platform for evaluating NET formation using HL-60-derived neutrophil-like cells (dHL-60). By employing GO as a representative nanomaterial, we demonstrated that dHL-60 cells recapitulate hallmark features of NETosis, including extracellular chromatin release, ROS production, and characteristic morphological changes, traditionally observed in primary neutrophils. Notably, GO induced NET formation in a concentration-dependent manner, and also suppressed PMA-induced NETosis, revealing its dual modulatory capacity and highlighting the complexity of nanomaterial-immune cell interactions. Our findings underscore the utility of dHL-60 cells as a viable alternative to primary neutrophils for immunotoxicological studies. The system provides consistent performance without the variability associated with donor-derived cells, and is amenable to high-throughput screening, mechanistic dissection, and standardization. Furthermore, the correlation between GO aggregation state and its bioactivity emphasizes the importance of physicochemical characterization in nanotoxicology assay design.

While the current work focused on GO, the assay platform is readily applicable to a broad spectrum of environmental, pharmaceutical, and industrial compounds known or suspected to modulate neutrophil function. Future studies incorporating structurally and mechanistically diverse substances will help to refine the system’s predictive value and expand its regulatory relevance. As regulatory frameworks shift toward mechanism-based and animal-free safety assessment strategies, the integration of NET-focused assays using standardized cell systems like dHL-60 holds promise for transforming the evaluation of immune-related toxicities across a wide array of chemical exposures.

